# The hippocampus supports interpolation into new states during category abstraction

**DOI:** 10.1101/2024.05.14.594185

**Authors:** Theo A.J. Schäfer, Mirko Thalmann, Eric Schulz, Christian F. Doeller, Stephanie Theves

## Abstract

The hippocampus forms concepts by integrating multi-feature relations into a unified representation. A common yet unconfirmed assumption is that such cognitive maps afford interpolations to never-experienced states. We approach this question as a category-learning problem in which prototypes are omitted from training but guide category-based decisions in a subsequent feature-inference task. Consistent with behavior, missing inferred stimulus features were represented at prototypical values in neocortex. This cortical completion effect correlated with hippocampal responses, which in turn reflected the distance between imagined prototypes and experienced exemplars. This was paralleled by a learning-dependent grid-like representation of the underlying conceptual space in entorhinal cortex. Our results suggest that abstracted prototypes correspond to interpolated central states in a cognitive map that guide cortical pattern completion during category-based decisions.

## Introduction

Human cognition fundamentally relies on a vast network of interconnected concepts that structure our experiences by facilitating the classification and differentiation of objects and events based on their shared and unique features. Prominent theories of concept learning posit that objects and events are represented in multidimensional psychological spaces (Gärdenfors, 2004; Nosofsky, 1984; Reed, 1972; Shepard, 1987). In these spaces, each experience can be portrayed as a coordinate with a particular combination of features, whereby categories correspond to regions. Which properties of psychological spaces are integral to concept representations during classification or inference has been debated for decades. A significant distinction concerns the extent to which idiosyncratic or abstract features are represented. Whereas exemplar theories propose the encoding of unique, experienced feature combinations, i.e. coordinates within psychological space (Kruschke, 1992; Medin & Schaffer, 1978; Nosofsky, 1984), another account suggests the formation of abstract prototypes, that reflect the central tendency of experiences (Homa et al., 1973; Posner & Keele, 1968; Reed, 1972; Rosch & Mervis, 1975). Mathematical models of both types of representations have gained empirical support at the behavioral level (Nosofsky, 1988; Smith & Minda, 2000). Especially coherent categories whose members share many features (high family resemblance) are thought to favor the formation of prototypes as an adaptive way to extract essential information and average out idiosyncratic noise (Minda & Smith, 2001; Smith et al., 2016). It is conceivable that different representational formats may coexist and adapt to different task demands. Accordingly, previous fMRI studies revealed that behavioral model estimates of prototype and exemplar representations correlate with activation levels in different brain regions (Bowman et al., 2020; Bowman & Zeithamova, 2018; Mack et al., 2013), whereby the hippocampal signal tracked model estimates of a prototype representation (Bowman & Zeithamova, 2018). Critically, while prototype representations are model assumptions derived from categorization behavior, it is unknown whether the brain actually represents the central tendency of experienced feature combinations. In fact, little is known about the neural representational basis of these estimates or the particular mechanisms by which the hippocampal formation might support prototype-based decisions.

It has been suggested that the representational schemes of the hippocampal-entorhinal system may critically support the encoding and retrieval of conceptual knowledge (Morton & Preston, 2021). Specifically, the system has been proposed to organize experiences and their relational properties into map-like representations (O’Keefe & Nadel, 1978; Tolman, 1948) based on an array of spatially-tuned cell types such as place (O’Keefe & Dostrovsky, 1971) and grid cells (Hafting et al., 2005), which encode positional and directional information (Bush et al., 2015; Moser et al., 2015). Similar mechanisms have been observed in encoding relations in spatial and non-spatial tasks (Aronov et al., 2017; Constantinescu et al., 2016; Garvert et al., 2017; Nau et al., 2018; Park et al., 2020; Tavares et al., 2015; Theves et al., 2019, 2020), suggesting domain-general representations (Behrens et al., 2018; Bellmund et al., 2018), considered beneficial for inference and generalization (Whittington et. al., 2020). Accordingly, these mechanisms seem to be influenced by behavioral relevance: for instance during concept learning, the hippocampus selectively encoded relations between exemplars along those feature dimensions that defined category membership (Theves et al., 2020). Importantly, while a commonly hypothesized feature of cognitive maps pertains to their metric function (Gärdenfors, 2004), allowing interpolations to non-experienced states, this property remains to be demonstrated for hippocampal processing.

Here we ask whether prototype representations inferred from behavior, manifest neurally as central states in a cognitive map. To this end, we use a category learning task in which the prototypes (i.e., the centroid feature combination per category space) are omitted during training, but guide category-based decisions in a subsequent feature inference task. In sum, we find evidence for pattern completion into central states of a hippocampal-entorhinal concept representation that guide cortical instatement of prototypical features during category-based decisions.

## Results

### Feature inference is anchored to category prototypes

In the present concept learning experiment (see Fig. 1A for an overview of the procedure), participants learned to categorize cartoon stimuli into three categories based on the combination of their two features (see Fig.1A-D) and then performed a feature inference task (Fig. 1E) in the MRI scanner. The category prototypes, denoting the centroids of the categories, were not shown during categorization training. Participants performed the categorization task until they reached 90% accuracy in the last two blocks or completed a maximum of 20 blocks. Participants’ responses indicate that they learned the category structure well: Categorization performance improved between the first and the last five blocks, both in terms of accuracy (*t*_46_ = 18.164, *P* < .0001) and response time (*t*_46_ = -5.600, *P* < .0001; see also Figure S1). On average, accuracy exceeded chance level (33 %) across the last five training blocks (*M* = 80.693 %, *t*_46_ = 41.578, *P* < .0001) and remained above chance in the final categorization test at the end of the experiment (*M* = 66.887 %, *t_46_* = 28.228, *P* < .0001).

**Figure 1:**
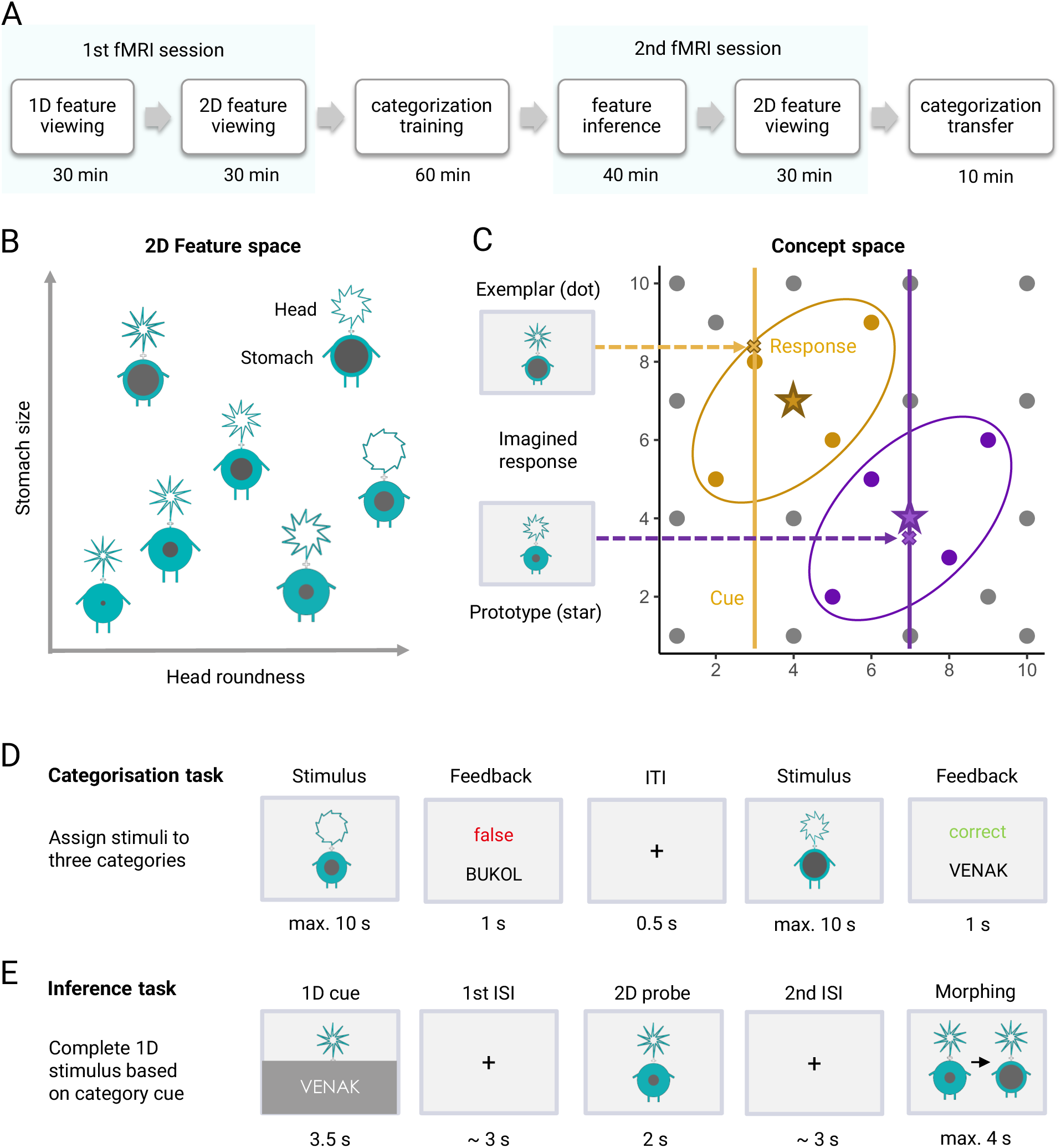
Experimental design and 2D feature space concept learning task. **A:** Overview of experimental tasks and sessions. In session 1, 1D and 2D feature viewing tasks (see Fig. S2) were followed by categorization training. In session 2, participants performed a feature inference task, a 2D feature viewing task, and a categorization test. **B:** Stimuli varied in the roundness of their heads and the size of their stomachs. **C** & **D:** Stimuli (dots) belonged to one of two elliptical-shaped categories (yellow & purple; labelled ‘Venak’ or ‘Bukol’) or to a residual category (gray). On each trial, participants assigned a stimulus to one of the three categories and received feedback. Training included only a subset of feature combinations, omitting the prototypes (stars). **E:** In the feature inference task, partial stimuli (e.g., consisting of the head) had to be completed by the occluded feature (e.g., the stomach) based on the category label. For illustration purposes, the stimulus size in figures D and E was increased, and the black background of the presentation screen changed to light gray.

In the subsequent feature inference task, participants were cued with a partial stimulus (including only one of the two features) and had to complete it by the missing second feature to generate a member of a given category. Specifically, they were instructed to imagine a potential category member with the cued feature and subsequently morph a probe stimulus, featuring a randomly sampled value of the previously omitted dimension, into the imagined one. We evaluated whether the completed feature was closer to the prototype or to the previously experienced cued exemplar by comparing the negative absolute distances to both locations (see Methods). We find that participants’ completion responses were closer to the prototype than to the cued exemplar (*t*_46_ = 6.404, *P* < .0001; Fig. 2), suggesting that feature inference was guided by an abstracted representation. The prototype bias in the completion responses was additionally confirmed by a comparison of model-based proximity scores (*t*_46_ = 9.527, *P* < .0001). Here, prototypes were defined as the means of multivariate Gaussian distributions, and the likelihood of a given coordinate within that distribution was converted into a proximity score. The scores were compared with the proximity scores of a Bayesian version of the Generalized Context Model (Nosofsky, 1984), which estimates the similarity of a stimulus to a category by the weighted sum of distances to all exemplars.

**Figure 2:**
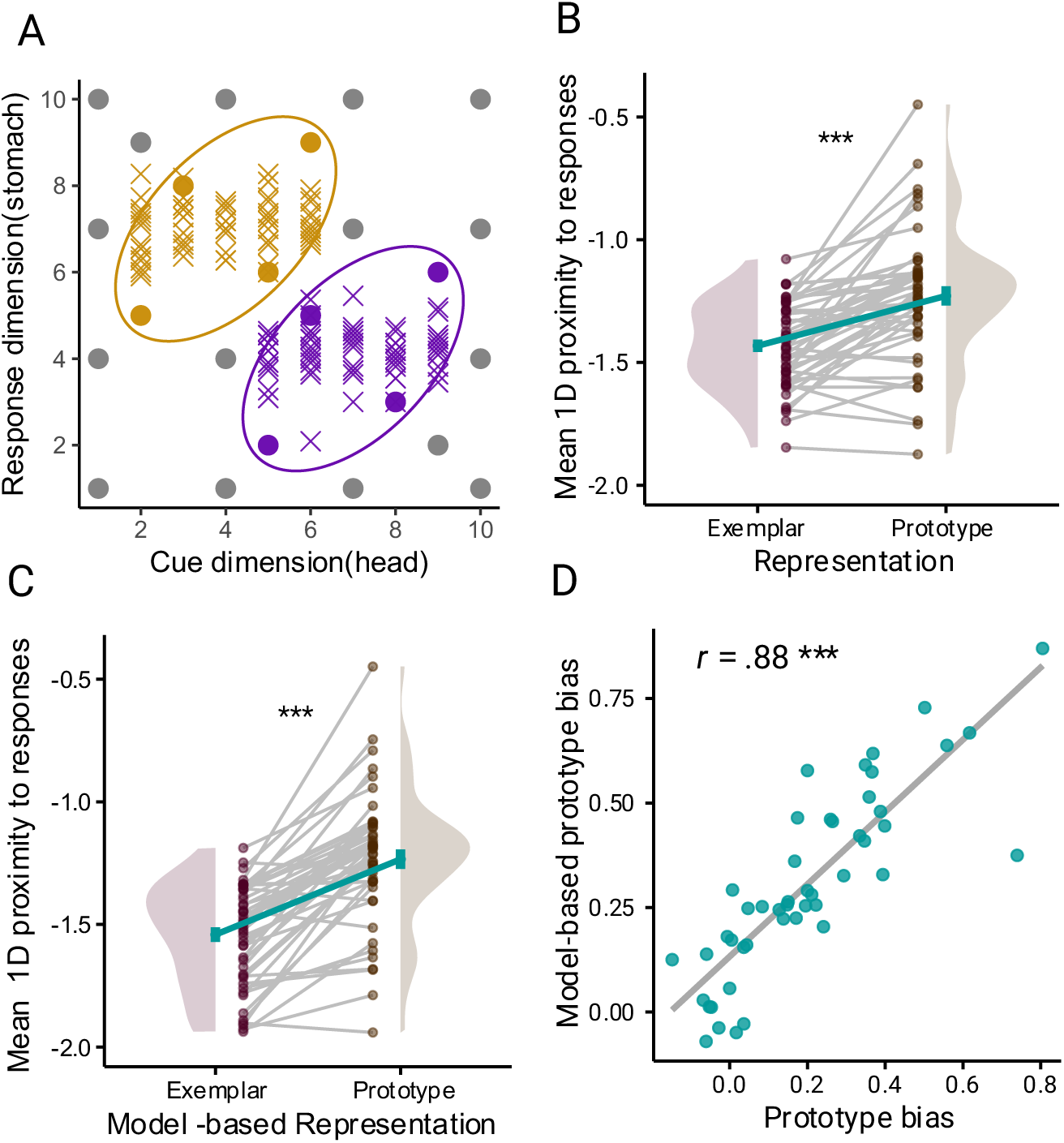
Behavioral completion bias indicates prototype representation. **A:** Completion responses (Xs) in the inference task of one example participant, cued on the ‘head’ dimension (x-axis) and responded on the ‘stomach’ dimension (y-axis). **B:** Proximity to the exemplar or prototype location is calculated as the negative absolute distance on the response dimension for each response and averaged per participant. Responses were closer to the prototype than to the cued exemplar (prototype bias). **C:** Proximity scores derived from Bayesian versions of prototype and exemplar models were based on the distance between predicted and participant’s completion responses. Model-based prototype proximity was higher than exemplar proximity. For B and C: Dots depict the proximity scores per participant for each condition; green lines with error bars correspond to means ± standard errors of the mean (SEM); distributions reflect probability density functions of data points. *** *P* < .001. **D:** Correlation between the measures in B and C. Green dots depict participants with a gray linear regression line. *** *P* < .001

### Grid-like representation of conceptual space in entorhinal cortex

First, we tested for a representation of the conceptual space by means of a grid-like representation in the entorhinal cortex. Grid cells in the entorhinal cortex fire at multiple locations of an environment in a regular hexagonal pattern (Hafting et al., 2005), with population dynamics providing a metric for space (Bush et al., 2015; Moser et al., 2015). FMRI proxies of grid-like activity in humans (i.e., directional modulation of the fMRI signal with 6-fold rotational symmetry) have been observed during transitions through physical and feature spaces (Doeller et al., 2010; Bao et al., 2019; Constantinescu et al., 2016; Nau et al., 2018; Viganò et al., 2021). For the purpose of our analysis, we treated stimulus successions in the 2D stimulus viewing blocks as trajectories of a given angle through conceptual space (Fig. 3A) and evaluated entorhinal pattern similarity between trajectory pairs as a function of their angular difference in 60°-space. Accordingly, our model representational dissimilarity matrix (RDM) predicts that the closer the angular difference between two trajectories is to multiples of 60°, the higher the similarity between the multivoxel- patterns elicited by those trajectories (i.e., highest similarity would be expected for trajectories multiples of 60° apart, and lowest similarity for trajectories multiples of 60°+30° apart). We find a significant correlation of the model RDM with entorhinal pattern similarity in the post-categorization stimulus viewing block (Fig. 3C bottom, *R* = .004, *t*_46_ = 2.034, *P* = .024). The effect was specific to a 6-fold rotational symmetry and was not present when 4-to-8-fold symmetries were used as controls (all *P* > .219, Fig 3C). As predicted, the effect was not present in the pre- categorization viewing block (Fig. 3C top, *R* = .001, *t*_46_ = .581, *P* = .282) when participants had not yet explicitly considered the relation of both features for categorization, consistent with the notion that cognitive map formation might be fostered by task demands (Theves et al., 2020). Finally, the strength of the grid-like feature space representation correlated with the behavioral prototype bias (Spearman’s rho = .31, *S* = 11870, *P* < .032, two-sided; correlation specific to 6-fold symmetry; other *P* > .15).

**Figure 3:**
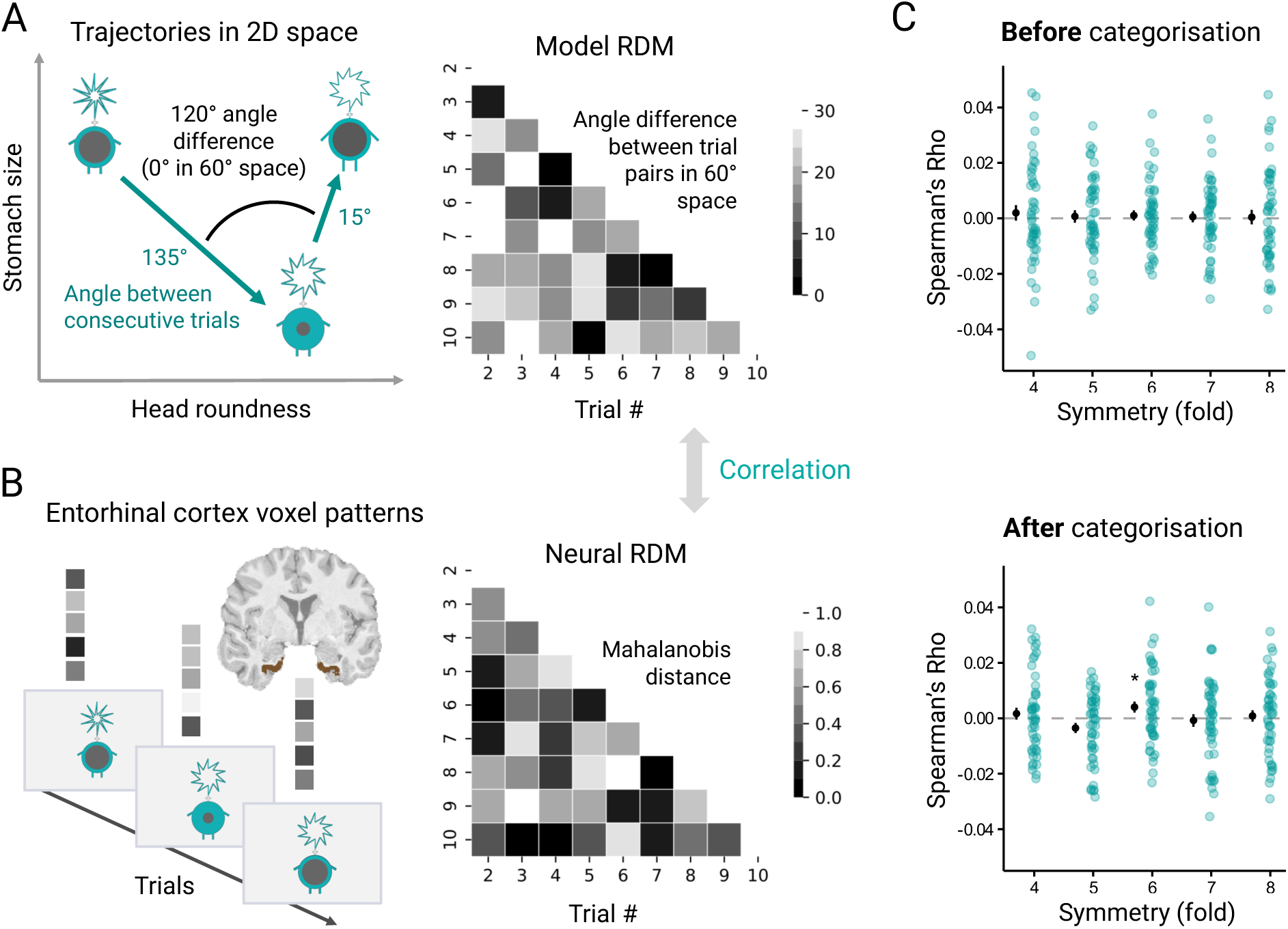
Grid-like representation of the conceptual space in entorhinal cortex. **A:** Logic of the hexadirectional analysis: Stimulus successions in the pre- and post-categorization 2D feature viewing tasks were treated as trajectories through feature space with a specific angle. The model RDM reflects the angular difference between trial pairs in 60° space (6-fold rotational symmetry; only a subset of trials is displayed in the RDMs). **B:** The neural RDM reflects the pairwise Mahalanobis’ distances between trial-wise fMRI BOLD activity estimates in voxels of an entorhinal cortex ROI. **C:** Run-wise correlation values (Spearman’s rho) between the model and neural RDMs were averaged for each participant and tested against zero in one-sample t-tests, for the pre- and post-categorization tasks and for control symmetries. Green dots refer to participant data points and black dots to the average correlation values +- SEM for each rotational symmetry. * *P* < .05.

### Pattern completion into prototypical features in visual cortex

Having established a behavioral prototype bias in the task as well as a representation of the conceptual space in the hippocampal-entorhinal system, we moved on to assess how prototype-based decisions are reflected in hippocampal processing. To evaluate whether feature inference was guided by neural exemplar or prototype representations, we applied a cross-task decoding approach in which we estimated the imagined feature value (e.g., the head) based on multi-voxel patterns in the imagination periods (empty screen) after the cue (e.g., the stomach) (Fig. 4A). We first trained and tested a support vector regression (SVR) algorithm to predict the values of the two single visual features on a given dimension based on visual cortex patterns during the 1D feature viewing task. A leave-one-run-out cross-validation procedure revealed significant above-chance decoding performance on both feature dimensions (head: mean Pearson correlation: *R* = .837, *t*_24_ = 44.140, *P* < .0001; negative mean absolute error = -1.414, *t*_24_ = 20.043, *P* < .0001; stomach: *R* = .843, *t*_21_ = 38.380, *P* < .0001; negative mean absolute error = -1.331, *t*_21_ = 20.218, *P* < .0001; Fig. S3 and S4). The validated decoder was then applied to voxel patterns in the imagination periods in the completion task to output continuous values on the missing feature dimension. We find that the decoded values in the second imagination period of the completion task were significantly close to the prototype, and significantly closer to the prototype than to the nearest exemplar (neural prototype bias: prototype proximity > exemplar proximity; *t*_46_ = 2.108, *P* = .020; proximity to the prototype: *t*_46_ = 4.619, *P* < .0001; proximity to the cued exemplar: *t*_46_ = 3.402, *P* = .001). This neural prototype bias correlated with the behavioral prototype bias (Spearman’s rho = .32, *S* = 11716, *P* = .014). There was no effect in the first imagination period (all *P* > .39). Importantly, the cortical completion into the prototypical features correlated positively with the cue-evoked hippocampal signal (Spearman’s rho = .27, *S* = 12682, *P* = .035; Fig. 4D), corresponding to fMRI signatures of pattern completion during episodic recollection (Horner et al., 2015). The higher hippocampal activity was in the post-cue period, the stronger was the subsequent instatement of cortical prototype-features. Interestingly, compared to previous studies on pattern completion, these correspond to an abstraction rather than an experienced event.

**Figure 4:**
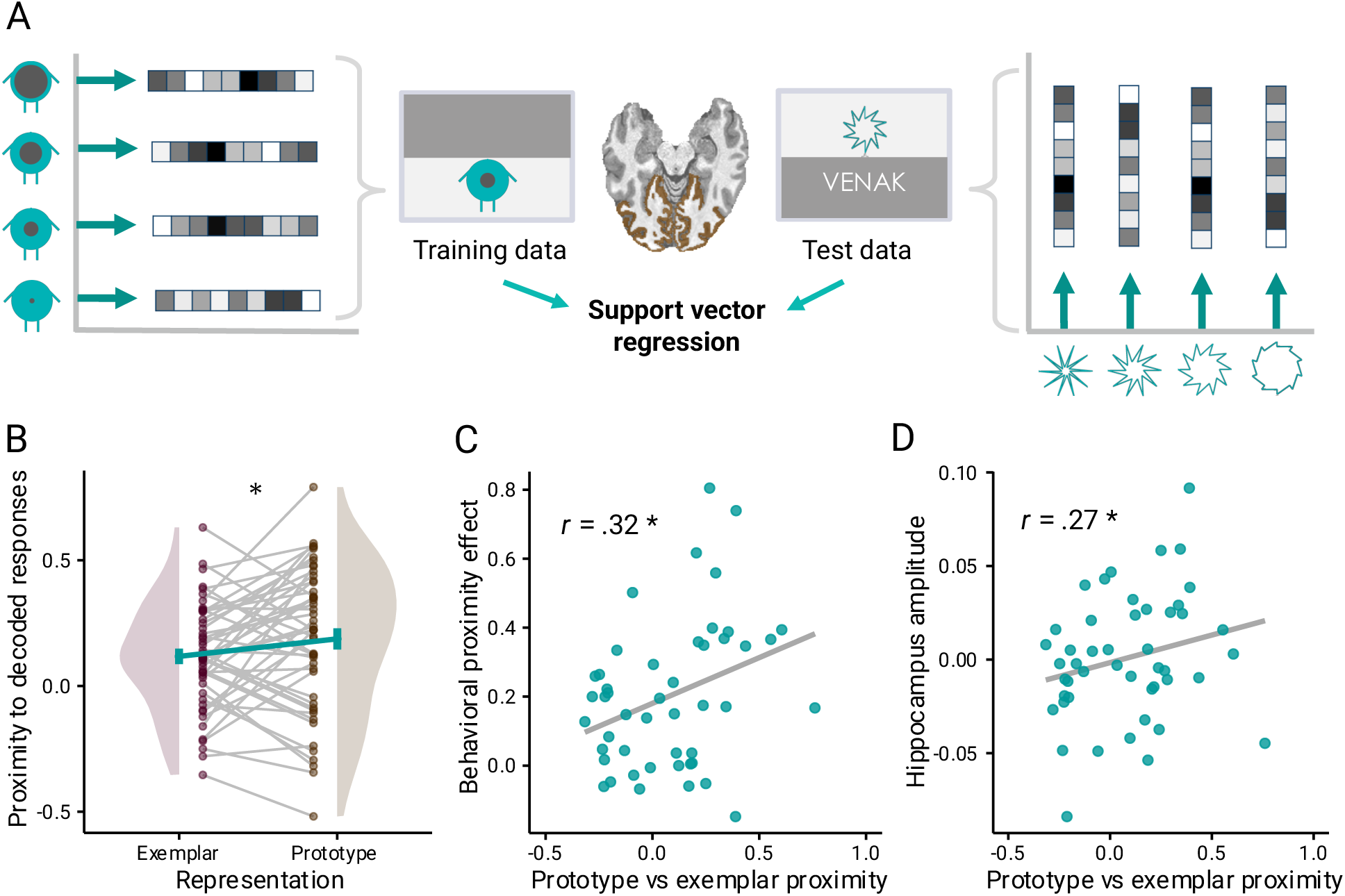
Instatement of prototypical features in visual cortex and correlation with behavior and hippocampal activation. **A:** The representation of the missing feature in visual cortex during the completion task was assessed via a cross-task decoding approach using support vector regression. After being trained and validated on the independent 1D-feature stimulus viewing task (Fig. 1A), the decoder was applied to voxel patterns in the post-cue interstimulus interval periods of the completion task. **B:** Decoded values were closer to the prototype than to the cued exemplar across participants (neural prototype bias). Dots depict the proximity scores per participant for each condition, green lines with error bars correspond to means ± SEM; distributions reflect probability density functions of data points. **C:** The neural prototype bias (x-axis) correlated significantly with the behavioral prototype bias (y-axis, see behavioral results in Fig. 2B) across participants. **D:** The neural prototype bias (x-axis) correlated with the cue-evoked mean amplitude in the hippocampus (y-axis) across participants. C and D: Green dots depict participants with a linear regression line in gray. * *P* < .05

### Hippocampal activation reflects representation of prototype position in conceptual space

The latter finding (Fig. 4D) might indicate that the hippocampus directs the retrieval of an unseen prototype representation, abstracted over experiences. Next, we assessed the representational content of the hippocampal signal that correlates with the cortical instatement of prototype features. To evaluate whether prototypes are incorporated into a cognitive map, we tested for a representation of the two-dimensional distance between prototype and surrounding exemplars. Specifically, we tested whether hippocampal adaptation during the probe stimuli scales with the two-dimensional distance of the probe stimulus to the prototype considered to be imagined in the preceding time window. Thus, we do not only evaluate the presence of a prototype representation, but test a model in which the prototype’s relation to exemplars corresponds to their distance in the two-dimensional conceptual space.

Indeed, we find a significant positive modulation of the BOLD response by prototype distance in the right hippocampus (cluster peak: *t*_46_ = 4.2, *P* = .016, [35, −21, −18]; subcluster peak: *t*_46_ = 3.44, *P* = .035, [33, −11, −24]; Figure 5B; Table S1 for an exploratory whole-brain analysis). There was no significant signal modulation by exemplar distance in the hippocampus. Furthermore, including both modulators in the same GLM yielded similar results, with clusters surviving only for prototype-distance modulation (cluster peak: *t*_46_ = 3.2, *P* = .019, [20, −11, −15] and *t*_46_ = 2.72, *P* = .038, [35, −21, −18]).

**Figure 5:**
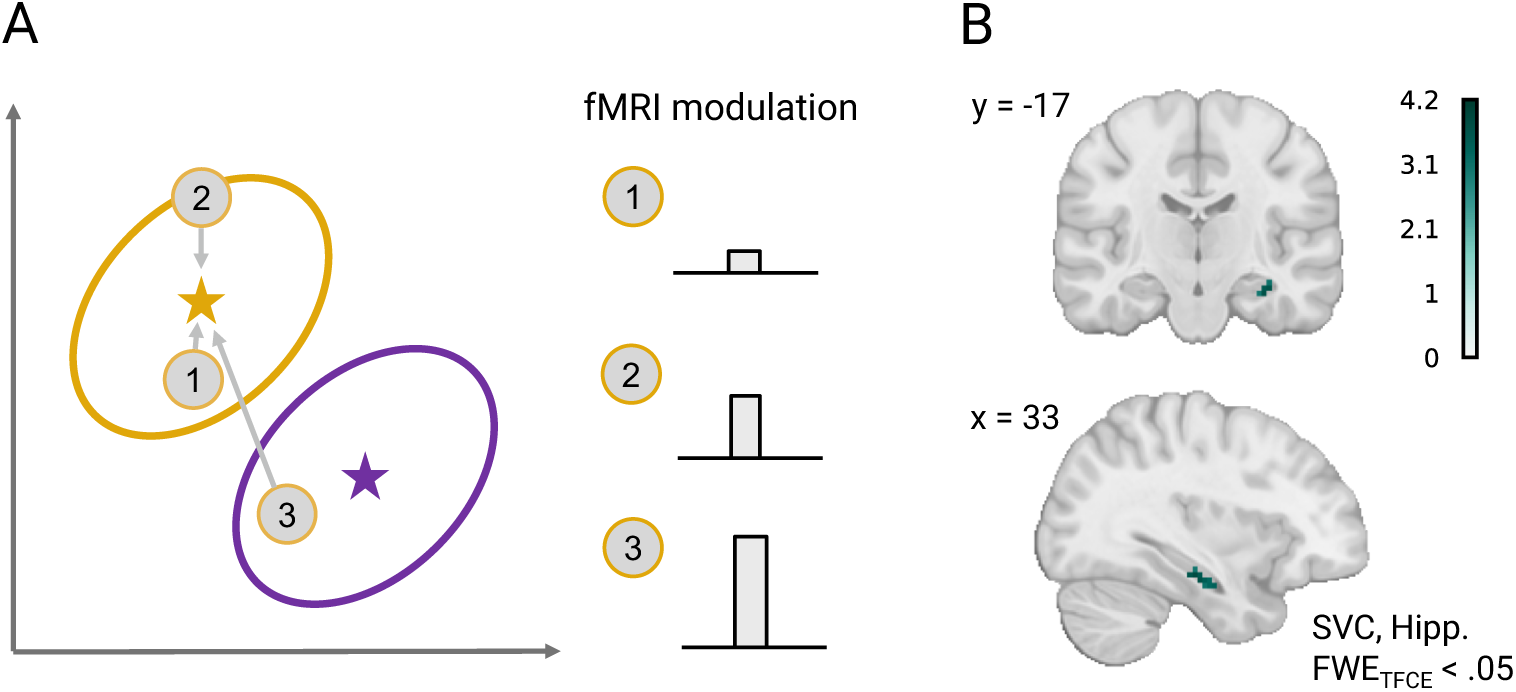
Hippocampal adaptation scales with probe stimulus’ 2D-distance to the prototype. **A:** Probe stimuli during the inference task (depicted as numbers) vary in their distance (arrows) to the prototype location (stars). Representational predictions were tested via fMRI adaptation analysis: If hippocampal representations of imagined prototype and experienced probe stimuli correspond to central and surrounding locations in a representational space, and if such a prototype representation is accessed in the post-cue period, a subsequent probe stimulus close to the prototype (1) should elicit a lower response than a distant probe stimulus (3). **B:** Supra-threshold (*P*_FWE_ < .05, TFCE; SVC) clusters in the right hippocampus that were modulated by the 2D distance to the prototype (displayed on the MNI template).

## Discussion

We report fMRI evidence for a neural correspondence to the cognitive-computational notion of categorical prototype representations. We find that, congruent with the emergence of an entorhinal grid-like representation of the underlying feature space, the hippocampus represented the distances between abstracted prototypes and presented exemplars. During feature inference, the hippocampal signal covaried with the instatement of prototypical values of the missing feature in visual cortex. Taken together, our findings suggest that the hippocampus represents prototypes as central states in a representational space that may guide pattern completion into neocortical representations during category-based decisions.

The present feature inference task allowed a continuous readout of representations from behavior and brain activity patterns and accordingly fine-grained comparisons of completion responses to different locations in feature space. Participants’ behavioral and neural responses were closer to the category prototype than to the cued exemplar. This prototype bias was also confirmed when comparing Bayesian versions of a prototype model and the Generalized Context Model (Nosofsky, 1986) which considers all members of a given category. Previous research demonstrated that the emergence of prototype representations depends on several factors, including the size of the training set (Minda & Smith, 2001), its coherence (Bowman & Zeithamova, 2020, 2023), and the amount of training (Smith & Minda, 1998; Bowman et al., 2020). These factors might explain differences in representation estimates across studies (Bowman & Zeithamova, 2018; Mack et al., 2013), and it is possible that exemplar biases could also emerge in the present task under different training conditions. However, the purpose of this study was to investigate the neural processes in a scenario in which behavior indicates prototype representations.

Previous studies have documented the involvement of the medial temporal lobe in concept learning (Bowman & Zeithamova, 2018; Davis et al., 2012; Kim et al., 2018; Kumaran et al., 2009; Liu et al., 2023; Nomura et al., 2007; Nomura & Reber, 2012; Schlichting et al., 2021; Seger et al., 2015; Zeithamova et al., 2008). Specifically, the hippocampus has been shown to encode relations between exemplars along category-defining feature dimensions (Theves et al., 2019, 2020), to readjust object representations to task-relevant features via attentional biases (Mack et al., 2016), and its activity amplitude covaried with prototype estimates during categorization (Bowman et al., 2020; Bowman & Zeithamova, 2018). Our findings significantly extend previous research by providing evidence for a representational mechanism by which the hippocampal system might support prototype abstraction. In particular, we observed a correlation between the hippocampal signal and neocortical representations of the missing features, similar to fMRI studies on pattern completion during episodic recollection (e.g. Horner et al., 2015; Grande et al., 2019). In contrast to previous research that focused on partial cue-evoked reinstatement of experienced event representations, the present findings might reflect hippocampal pattern completion (for review, see Theves et al., 2024) into previously unseen feature combinations (the prototypes) that can be abstracted from experiences. Consistent with a potential pattern completion account, the representation of the missing prototype-proximal feature in visual cortex followed a hippocampal representation of the prototype location and reached significance in the second, not the first ISI. The hippocampal representation of the prototype location was evidenced by signal adaptations reflecting the distances between the prototype and surrounding exemplars in conceptual space. Hippocampal pattern completion can be influenced by attractor states that are formed by recurrent connections within CA3 of the hippocampus (McNaughton & Morris, 1987; Treves & Rolls, 1992). Patterns close to an attractor will settle into the attractor state. In this vein, neural and behavioral completion biases could result from an integration of cued category members into attractors corresponding to the prototypical states.

Concurrently, we observed a grid-like representation of the underlying feature space in the entorhinal cortex, which provides the major input to the hippocampus. Grid cell firing is assumed to reflect the latent structure of an environment (Stachenfeld et al., 2017; Whittington et al., 2020; Spens & Burgess, 2024), which can support vector-based navigation and structural generalization more broadly. A recent memory model (Spens & Burgess, 2024) further suggests that shared category features might initially be stored in entorhinal cortex as latent variables that are used for memory retrieval in the hippocampus. In the present study, the grid-like representation was present only after, not before, participants completed the categorization training. This is in line with a previous finding that suggests that the cognitive demand of integrating both feature dimensions, e.g. to assign category membership, fosters hippocampal concept representations (Theves et al., 2020). The strength of the entorhinal grid-like representation further correlated with participants’ behavioral prototype bias. Future investigations could explore entorhinal representations of less cohesive categories (e.g., with highly heterogeneous exemplars or including exceptions), that typically favor exemplar representations (Bowman & Zeithamova, 2020, 2023; Minda & Smith, 2001).

In sum, the present findings advance our understanding of the nature of concept representations by suggesting a neural mechanism underlying the behaviorally-inferred use of category prototypes. We show that prototype-guided behavior during category-based feature inference is accompanied by neural patterns that reflect central positions or interpolations with respect to the representation of other exemplars. On a more general level, this finding might relate hippocampal processing to a commonly assumed property of cognitive maps: The implicit representation of “what lies between” experienced states, which allows the interpolation to not directly experienced states. Such a property is pivotal to the idea that cognitive maps endow flexibility to cognitive operations, for instance in imagination, planning, or the formation of task-efficient representations like prototypes.

## Materials & Methods

### Participants

Fifty volunteers participated in the study. Participants indicated right-handedness, no present or previous neurological or psychiatric disease and normal or corrected-to-normal vision. One participant was excluded because fMRI data acquisition and the task were interrupted several times due to technical problems, leading to an incomplete dataset. Two participants were excluded, because we noticed inconsistencies in the stimulus timing during data collection and preprocessing. Thus 47 participants were included in the analyses (age: *M* = 27.4, *SD* = 4.34, range = 18-35; gender: male = 23, female = 24). All participants gave informed written consent prior to participation and were compensated for participation. The study was approved by the ethics committee at the Medical Faculty of the University of Leipzig.

### Experimental Procedure

The experiment consisted of several tasks performed over the course of one day (see Figure 1A). In a behavioral session, participants learned to categorize visual stimuli based on the combination of two continuous features. Subsequently, they performed a feature inference task in the MRI scanner, in which partial category stimuli had to be completed by the missing feature. The concept learning and inference tasks were preceded and followed by stimulus viewing blocks in the MRI scanner. There was a 1-hour break between the category learning task and the inference task.

### Stimuli

Cartoon figures were used as stimuli across all tasks. These figures varied along two continuous dimensions: the size of their stomach and the roundness of their head (Figure 1B). Stomach size was defined as the radius of the grey-filled circle inside the turquoise body, while roundness of the head was defined as the radial distance between two circles’ inner and outer vertices, which together formed a head with ten spikes. Head and body were connected by a white fixation cross and the figures were presented against a black background. In the 1D feature viewing and the feature completion task, half of the display was covered by a dark gray rectangle. The stimuli were projected onto a screen via a mirror attached to the MRI head coil. Participants responded using either a computer keyboard outside the scanner or MRI-compatible button boxes inside the scanner. PsychoPy (version 2020.10.2) and custom Python (3.9.7) scripts controlled the generation and presentation of the stimuli.

### Experimental tasks

#### Categorization task

Category learning proceeded in a feedback-based classification task. Stimuli belonged to one of two elliptical-shaped categories located above and below the diagonal in a two-dimensional feature space, or to a residual category that served as their outer boundary (Fig. 1C). The purpose of the third category was to avoid competition between representations of the category centroids and the most extreme values of the feature space (“caricatypes”). On each trial, participants had to assign a stimulus to one of the categories (labeled VENAK, BUKOL or NONE) based on the combination of its features (Fig. 1D). Responses (corresponding to “v”, “b”, “n” on the keyboard) were followed by 1 s of corrective feedback (”correct”, “false”, or “too slow” after 10 s of no response). Trials were separated by a fixation cross for 0.5 s. Only a subset of the possible feature-value combinations was presented during training, the category centroids (prototypes) were omitted. Four stimuli were drawn from each of the two elliptic categories and repeated four times, while sixteen stimuli were drawn from the residual category and repeated one, so that all categories occurred with equal probability. After a training block in which participants chose between the two elliptic categories, they completed at least 5 blocks of 48 trials each in which they chose between all three categories. After each block, they received feedback on their performance (percentage correct). Training continued until a maximum of 20 blocks or 90 % accuracy across the last two blocks was reached.

#### Feature completion task

In a subsequent completion task inside the MRI scanner, participants were cued with a partial stimulus that had to be completed by the missing feature to generate a member of a given category (Fig. 1E). Specifically, one of the features (head or stomach, counterbalanced across participants) was occluded by a gray rectangle, and the first letter of the category label (”V” or “B”) was presented in white above or below the stimulus, respectively. Participants were instructed to imagine a member of the given category with the cued feature. Cue presentation (3.5 s) was followed by a fixation cross (ISI 1: average 3 s, sampled from a truncated exponential distribution with min = 2 s, max = 8 s, mu = 3 s), and then by a probe stimulus (2 s) consisting of the cued feature and a randomly sampled feature from the previously occluded dimension. The probe was followed by a second fixation cross (ISI 2), before it reappeared and was morphed by the participant into the imagined 2D stimulus. Participants increased or decreased the feature values using two buttons (right index and middle finger; a total of 100 steps were possible) and confirmed their choice with a third button press (left index finger). They had four seconds to respond before the trial ended. Five different cues containing all values for each elliptic category were presented, each repeated five times in each of three runs.

#### Feature viewing and reconstruction tasks

Participants performed two feature viewing tasks at the beginning of the experiment (1D stimuli, 2D stimuli) and one feature viewing task (2D stimuli) at the end of the experiment (Supplementary Figure S2). The blocks served to train feature decoders to be applied to the subsequent completion task and to investigate hexadirectional representations of the feature space as a function of category learning. In the 2D viewing task, participants were instructed to attend carefully to both features of the (complete) stimuli. Stimuli were presented for 2 s and separated by a fixation cross (intertrial interval: sampled from a truncated exponential distribution with min = 2 s, max = 8 s, mu = 3 s). Fifty-two feature-value combinations were presented, entailing five repetitions per feature value per dimension. Each 2D stimulus was repeated once in each of 4 runs. To ensure that participants attended to the stimulus values, we included 22 % test trials in each run. Test trials were indicated by a purple fixation cross (1 s) in between the two to be compared stimuli. In test trials, the stimuli had changed in one of the features and had to be reconstructed (response buttons and timing equivalent to the feature completion task). Feedback (0.5 s) in form of a green number indicated the distance in steps between their provided and correct response. After each run, participants received performance feedback, indicated by the average distance between their responses and the true feature value. Test trials were pseudo-randomly distributed, with the same number of test trials in each bin of 16 trials. The 1D stimulus viewing task was similar, except that one of the features (counterbalanced across participants) was occluded by a gray rectangle. Each of the 10 values from the presented dimension was shown 5 times in each of the 4 runs, together with 15 test trials (23 %).

#### Categorization test

At the end of the experiment, participants performed a categorization test outside the scanner in which no feedback was provided and stimuli from the entire feature space were presented twice for a total of 200 trials. In a final task, participants were asked to generate the most typical member of each category by reconstructing both features with two sliders. They completed two trials for each elliptic category.

#### Behavioral analysis

We analyzed categorization training data by averaging categorization accuracy across blocks and participants. To examine learning effects on performance, we compared the first and last five blocks in a paired t-test. Categorization performance at the end of the training was compared to chance level in a one-sided, one-sample t-test. Data of the feature inference task was analyzed with respect to the proximity of participants’ completion responses to the prototype and exemplar coordinates. Proximity was defined as the negative absolute distance between two values on the response dimension. Specifically, trial-wise completion values were compared to the centroids of the respective cued category (proximity to prototype) and to the exemplar location corresponding to the cue (proximity to exemplar). Considering that the feature-inference task entailed explicit cues, we considered the cued-/nearest exemplar the most intuitive comparison to the prototype prediction. For the cue corresponding to the prototype value, the maximal proximity to the neighboring exemplars was chosen. Mean proximities to prototype and exemplar locations per participant were compared in a one-sided, paired t-test. Analyses were performed in Python 3.9.7 using the Spyder developer environment (version 5.1.5) and in R 4.2.2 using the RStudio developer environment (version 2023.06.0). Custom Python scripts used the numpy, pandas, matplotlib, seaborn and scipy packages. R scripts used the tidyverse and ggplot2 packages. Unless otherwise noted, we tested directed hypothesis in one-sided tests at alpha level of 5 %.

#### Modeling concept representations

For comparability to studies that compared prototype estimates to the weighted sum of distances to all exemplars, we confirmed the observed prototype bias using the following model-based proximity scores. Prototypes were defined as the means of multivariate gaussian distributions and the likelihood of a given coordinate within the distribution was converted to a proximity score. The exemplar model was a Bayesian version of the Generalized Context Model (GCM, Nosofsky, 1984), which estimates the similarity of a stimulus to a category by the weighted sum of distances to all the individual training exemplars. We used the Bayesian statistical software RStan, a package for R that facilitates the estimation of Bayesian statistical models using Hamiltonian Monte Carlo (HMC). We defined the structure of the prototype and exemplar models in Stan’s probabilistic programming language. This involved specifying prior distributions for each model parameter (e.g., the location of the prototypes in feature space, the generalization parameter; for specification of priors, see https://github.com/MirkoTh/hierarchical-categorization). We fitted these models to the data using RStan’s sampling function, which draws from the posterior distribution of the model parameters given the data. The convergence of the models was checked using the Rhat statistic, with values close to 1 indicating good convergence. Effective Sample Sizes (ESS) were evaluated to ensure that the Markov chains had explored the parameter space thoroughly. The posterior distributions of the model parameters were analyzed, and point estimates were extracted as the maximum a-posteriori estimates (MAP). Based on the individually fitted parameters, we computed for each cue the most likely completion response based on each model, and calculated the proximity to the participant’s responses.

#### MRI data acquisition

MRI data were recorded using a 32-channel head coil on a 3 Tesla Siemens Magnetom SkyraFit system (Siemens, Erlangen, Germany). FMRI scans were acquired in axial orientation using T2*- weighted whole-brain gradient-echo echo planar imaging (GE-EPI) with multi-band acceleration, sensitive to blood-oxygen-level-dependent (BOLD) contrast (Feinberg et al., 2010; Moeller et al., 2010). The fMRI sequence had the following parameters: TR = 1500 ms, TE = 22 ms, voxel size = 2.5 mm isotropic, FOV = 204 mm, flip angle = 80°, partial Fourier factor = 6/8, bandwidth = 1794 Hz/Px, 63 interleaved slices, distance factor = 10 %, phase encoding direction = A-P, multi-band acceleration factor = 3. Field maps using the opposite phase-encoded EPIs were recorded between the task runs (Parameters: TR = 8000 ms; TE = 50 ms; voxel size = 2.5 mm isotropic; field of view = 204 mm; flip angle = 90°; partial Fourier factor = 6/8; bandwidth = 1794 Hz/Px; multi- band acceleration factor = 1; 63 slices interleaved; slice thickness = 2.5 mm; distance factor = 10 %) to correct for magnetic field inhomogeneities (see preprocessing below). At the end of the second scanning session, we acquired a T1-weighted MP2RAGE anatomical scan (TR = 5000 ms; TE = 2.9 ms; TI_1_ = 700 ms; TI_2_ = 2500 ms; voxel size = 1 mm isotropic; field of view = 256 mm; flip angle_1_ = 4°; flip angle_2_ = 5°; bandwidth = 240 Hz/Px; acceleration factor = 3; distance factor = 50 %). Participants were presented with task stimuli on a screen, which was viewed through a mirror attached to the head coil. Behavioral responses were recorded using MRI-compatible button boxes.

#### MRI preprocessing

Results included in this manuscript come from preprocessing performed using *fMRIPrep* 21.0.1 (Esteban, Markiewicz, et al. (2018); Esteban, Blair, et al. (2018); RRID:SCR_016216), which is based on *Nipype* 1.6.1 (K. Gorgolewski et al. (2011); K. J. Gorgolewski et al. (2018); RRID:SCR_002502).

### Preprocessing of B_0_ inhomogeneity mappings

A total of 2 fieldmaps were found available within the input BIDS structure. A *B_0_*-nonuniformity map (or *fieldmap*) was estimated based on two (or more) echo-planar imaging (EPI) references with topup (Andersson, Skare, and Ashburner (2003); FSL 6.0.5.1:57b01774).

### Anatomical data preprocessing

A total of 2 T1-weighted (T1w) images were found within the input BIDS dataset. All of them were corrected for intensity non-uniformity (INU) with `N4BiasFieldCorrection` (Tustison et al. 2010), distributed with ANTs 2.3.3 (Avants et al. 2008, RRID:SCR_004757). The T1w-reference was then skull-stripped with a *Nipype* implementation of the `antsBrainExtraction.sh` workflow (from ANTs), using OASIS30ANTs as target template. Brain tissue segmentation of cerebrospinal fluid (CSF), white-matter (WM) and gray-matter (GM) was performed on the brain-extracted T1w using `fast` (FSL 6.0.5.1:57b01774, RRID:SCR_002823, Zhang, Brady, and Smith 2001). A T1w-reference map was computed after registration of 2 T1w images (after INU-correction) using `mri_robust_template` (FreeSurfer 6.0.1, Reuter, Rosas, and Fischl 2010). Brain surfaces were reconstructed using `recon-all` (FreeSurfer 6.0.1, RRID:SCR_001847, Dale, Fischl, and Sereno 1999), and the brain mask estimated previously was refined with a custom variation of the method to reconcile ANTs-derived and FreeSurfer-derived segmentations of the cortical gray-matter of Mindboggle (RRID:SCR_002438, Klein et al. 2017). Volume-based spatial normalization to one standard space (MNI152NLin2009cAsym) was performed through nonlinear registration with `antsRegistration` (ANTs 2.3.3), using brain-extracted versions of both T1w reference and the T1w template. The following template was selected for spatial normalization: *ICBM 152 Nonlinear Asymmetrical template version 2009c* [Fonov et al. (2009), RRID:SCR_008796; TemplateFlow ID: MNI152NLin2009cAsym].

### Functional data preprocessing

For each of the 15 BOLD runs found per subject (across all tasks and sessions), the following preprocessing was performed. First, a reference volume and its skull-stripped version were generated using a custom methodology of *fMRIPrep*. Head-motion parameters with respect to the BOLD reference (transformation matrices, and six corresponding rotation and translation parameters) are estimated before any spatiotemporal filtering using `mcflirt` (FSL 6.0.5.1:57b01774, Jenkinson et al. 2002). The estimated *fieldmap* was then aligned with rigid-registration to the target EPI (echo-planar imaging) reference run. The field coefficients were mapped on to the reference EPI using the transform. BOLD runs were slice-time corrected to 0.705s (0.5 of slice acquisition range 0s-1.41s) using `3dTshift` from AFNI (Cox and Hyde 1997, RRID:SCR_005927). The BOLD reference was then co-registered to the T1w reference using `bbregister` (FreeSurfer) which implements boundary-based registration (Greve and Fischl 2009). Co-registration was configured with six degrees of freedom. Several confounding time-series were calculated based on the *preprocessed BOLD*: framewise displacement (FD), DVARS and three region-wise global signals. FD was computed using two formulations following Power (absolute sum of relative motions, Power et al. (2014)) and Jenkinson (relative root mean square displacement between affines, Jenkinson et al. (2002)). FD and DVARS are calculated for each functional run, both using their implementations in *Nipype* (following the definitions by Power et al. 2014). The three global signals are extracted within the CSF, the WM, and the whole-brain masks. Additionally, a set of physiological regressors were extracted to allow for component-based noise correction (*CompCor*, Behzadi et al. 2007). Principal components are estimated after high-pass filtering the *preprocessed BOLD* time-series (using a discrete cosine filter with 128s cut- off) for the two *CompCor* variants: temporal (tCompCor) and anatomical (aCompCor). tCompCor components are then calculated from the top 2% variable voxels within the brain mask. For aCompCor, three probabilistic masks (CSF, WM and combined CSF+WM) are generated in anatomical space. The implementation differs from that of Behzadi et al. in that instead of eroding the masks by 2 pixels on BOLD space, the aCompCor masks are subtracted a mask of pixels that likely contain a volume fraction of GM. This mask is obtained by dilating a GM mask extracted from the FreeSurfer’s *aseg* segmentation, and it ensures components are not extracted from voxels containing a minimal fraction of GM. Finally, these masks are resampled into BOLD space and binarized by thresholding at 0.99 (as in the original implementation). Components are also calculated separately within the WM and CSF masks. For each CompCor decomposition, the *k* components with the largest singular values are retained, such that the retained components’ time series are sufficient to explain 50 percent of variance across the nuisance mask (CSF, WM, combined, or temporal). The remaining components are dropped from consideration. The head-motion estimates calculated in the correction step were also placed within the corresponding confounds file. The confound time series derived from head motion estimates and global signals were expanded with the inclusion of temporal derivatives and quadratic terms for each (Satterthwaite et al. 2013). Frames that exceeded a threshold of 0.5 mm FD or 1.5 standardised DVARS were annotated as motion outliers. The BOLD time-series were resampled into standard space, generating a *preprocessed BOLD run in MNI152NLin2009cAsym space*. First, a reference volume and its skull-stripped version were generated using a custom methodology of *fMRIPrep*. The BOLD time-series were resampled onto the following surfaces (FreeSurfer reconstruction nomenclature): *fsnative*, *fsaverage*. All resamplings can be performed with *a single interpolation step* by composing all the pertinent transformations (i.e. head-motion transform matrices, susceptibility distortion correction when available, and co-registrations to anatomical and output spaces). Gridded (volumetric) resamplings were performed using `antsApplyTransforms` (ANTs), configured with Lanczos interpolation to minimize the smoothing effects of other kernels (Lanczos 1964). Non-gridded (surface) resamplings were performed using `mri_vol2surf` (FreeSurfer).

### fMRI data analysis

fMRI data was analyzed using Python 3.9.7 with the Spyder developer environment (version 5.1.5) and R 4.2.2 using the RStudio developer environment (version 2023.06.0). Custom Python scripts relied on the nilearn, nltools, sklearn, scipy, and rsatoolbox packages. R scripts used the tidyverse and ggplot2 packages. Correlations were computed using the Pearson correlation coefficient whenever both variables did not violate data normality according to the Shapiro-Wilk test. Otherwise, Spearman’s rank correlation coefficient (rho) was used as the correlation measure. Almost all our hypotheses on fMRI effects were clearly directed, i.e. based on previous literature. Thus, we used one-sided tests with an alpha level of 5%, unless otherwise noted.

### General information about first-level general linear models (GLMs)

First-level GLMs were implemented using the FirstLevelModel class of the nilearn Python package and computed within a brain mask in participants’ native space. The mask was based on the anatomical brain mask in native space created during preprocessing with fMRIPrep and resampled to the resolution of the functional data. Task-related regressors in the GLMs were convolved with the Glover haemodynamic response function (HRF). Temporal autocorrelation was accounted for using an autoregressive AR(1) model. Nuissance regressors included 24 motion regressors (3 translations, 3 rotations, their squares, their derivatives, and their squared derivatives), anatomical-component based noise correction components (aCompCor) regressors derived from fMRIPrep up to a sum of 15 % explanation of variance, 4 global signal regressors (including the square, derivative, and squared derivative), and discrete cosine-basis regressors estimated by fMRIPrep to account for low-frequency temporal drifts.

### Representational similarity analysis (RSA) of hexadirectional signal

To test for a grid-like representation of the feature space, we pursued a model-based RSA of the feature viewing task data. This analysis approach provided an efficient alternative to an active sampling of trajectory angles via an additional task (cf., Bao et al. 2019; Bellmund et al. 2016; Vigano et al. 2021, 2023). For this purpose, we treated stimulus successions in the 2D feature viewing tasks as trajectories of a given angle in conceptual space and analyzed the pattern similarity between trajectories as a function of their angular difference in 60° space via representational similarity analysis (RSA, Kriegeskorte et al., 2008). (In the task, frequent catch trials asked participants to morph a stimulus into the preceding one, making a general consideration of the vector between successive stimuli conceivable). BOLD fMRI data from the 2D feature viewing task was modelled in single-trial GLMs per run. Following the least-squares-separate approach (Mumford et al., 2012), each GLM included an onset regressor for one of the 52 feature combinations, an onset regressor for the remaining stimuli and additional task event regressors (onset of test stimulus, probe, response, and feedback), resulting in 4 (runs) x 52 parameter estimate (PE) maps. PE images were masked with participant-specific bilateral masks of the entorhinal cortex created by FreeSurfer segmentations of the participants’ anatomical images during preprocessing with fMRIPrep (FreeSurfer labels 1006 & 2006). The masked images were z-scored across conditions within each run and their pairwise Mahalanobis’ distances computed to generate a neural representational dissimilarity matrix (RDM). The model RDM was based on the difference in trajectory angle between each pair of trials in 60° space. The similarity between neural and model RDMs was estimated via Spearman’s rho, averaged for each participant, and the resulting correlation values were tested against zero in a one-sided, one-sample t-test. The specificity of a 6-fold modulation of activity was evaluated via control analyses with RDMs based on 4-, 5-, 7- and 8-fold rotational symmetries.

### Feature decoding analysis

To evaluate whether completion responses are guided by exemplar or prototype representations, we applied a cross-task decoding approach, in which values of the missing feature in the inference task were predicted based on a decoder that was trained on the preceding 1D feature viewing task. We trained a linear support vector regression (SVR) algorithm to predict feature values based on multi-voxel patterns during the 1D feature viewing task. The GLM of the 1D feature viewing task included an onset regressor for each of the 10 feature values and onset regressors for test events (test stimulus, probe, response and feedback), resulting in 4 (runs) x 10 different parameter estimate (PE) maps. The PE values served as training data for the decoder. For the test data, we modelled the feature completion task with a GLM that contained one regressor for each of the 10 cues (1D feature + category label) with onset and duration of the subsequent fixation cross and onset regressors for task events (the 2D probe, test onset and response onset), resulting in 3 (runs) x 10 (cues) PE maps. The PE maps were z-scored within each run and masked with participant-specific bilateral visual cortex masks created by FreeSurfer segmentations of the participants’ anatomical images during preprocessing with fMRIPrep. The mask included the pericalcarine cortex (FreeSurfer labels 1021 & 2021), cuneus (1005, 2005), lingual cortex (1013, 2013) and lateral occipital cortex (1011, 2011). A linear SVR decoder (with parameters c = 1 and epsilon = 0.1) was trained and validated on stimulus values and the multi- voxel features from the training GLM and then applied to predict stimulus values based on the multi-voxel features of the testing GLM. The proximity of the predicted values to either the cued exemplar or prototype was computed as negative absolute distance, akin to the behavioral analysis. Proximity values were compared to chance-level performance of the decoder by repeating the analysis 5000 times and randomly permuting the condition labels of the training data. The z-score of the actual proximity value within this permutation distribution was calculated for each subject. We tested prototype and exemplar proximity against zero in a one-sided, one-sample t-test and their difference in a one-sided, paired two-sample t-test. We applied this analysis to both ISI periods. We initially validated the decoder performance on the training data in a leave-one-run-out SVR analysis. Decoding performance was measured by the negative mean absolute error as well as by the Pearson correlation coefficient between the decoded values from each of the 4 runs and the true stimulus values. Participant’s averages of these measures were tested against a permutation derived chance level in a one-sided, one-sample t-test. The neural prototype bias was correlated with the behavioral prototype bias and the hippocampal signal during the post-cue period. For the latter, we averaged the PE maps of the post-cue imagination period from the feature completion task GLM across runs and across voxels within a hippocampal mask (FreeSurfer labels: 17 & 53). This resulted in a hippocampal signal value for each participant which was correlated with the individual neural prototype bias values.

### Adaptation analysis

We evaluated prototype representations during the feature inference task using fMRI adaptation analysis. Retrieval of the prototype after the cue would be reflected in lower responses to subsequent probe stimuli, the closer they are to the prototype location. The GLM included onset regressors for the cue stimuli, the probe stimulus, the morphing phase and the response. The probe stimulus regressor was accompanied by a parametrically modulated regressor (demeaned) denoting the Euclidean distance between the probe and the prototype location. For control analyses, we ran a GLM with a parametric regressor reflecting the distance to the cued exemplar, as well as a GLM including both parametric regressors. PE images were transformed to MNI standard space using ANTS (version 2.3.5), resampled to the resolution of the functional data, and spatially smoothed with a 7.5 mm full width at half maximum Gaussian filter (FWHM). For group-level statistics, we performed a mass-univariate nonparametric test (Freedman & Lane, 1983) with 5000 permutations using threshold-free cluster enhancement (TFCE) within a small volume correction (SVC) hippocampal mask (FreeSurfer labels 17 & 53) and corrected for multiple comparisons with a family-wise error rate (*P*_FWE_ < .05). Labels of significant brain clusters were extracted via the Harvard-Oxford (Sub)Cortical Structural Atlas.

## Acknowledgements

We thank Alexander Nitsch and Felix Deilmann for jointly writing Python scripts for MRI task presentation and data analyses, and Ulrike Horn for contributing code for operating the MRI button box. We thank Kerstin Schumer, Anke Kummer, Simone Wipper, Sylvie Neubert, Mandy Jochemko, Nicole Pampus and Manuela Hofmann for their assistance in data collection. We thank Rebekka Tenderra for initially piloting the grid analysis approach on data from Theves et al, 2019. We thank the University of Minnesota Center for Magnetic Resonance Research for providing the multiband EPI sequence software and Toralf Mildner, Joeran Lepsien and colleagues of the Psychology Department and the Adaptive Memory research group for cooperating in MRI sequence piloting, as well as colleagues of the Psychology Department for discussions regarding the study. ST’s research is supported by a Minerva Fast Track fellowship of the Max Planck Society. CD’s research is supported by the Max Planck Society, the European Research Council (ERC-CoG GEOCOG 724836), the Kavli Foundation, the Jebsen Foundation and Helse Midt Norge.

## Author contributions

ST conceived and designed the experiment. TS and ST designed and prepared the tasks. TS programmed the tasks and acquired the data. TS and ST analyzed the data. MT performed the model-based behavioral analysis with input from ES. TS wrote the initial draft. TS and ST jointly wrote the manuscript. CD provided general advice and contributed to the manuscript. All authors discussed the project.

## Data availability

Data to reproduce the statistical analyses reported in this paper will be made available upon publication via the Open Science Framework.

## Code availability

The analysis code will be available upon publication on GitHub.

## Supplemental material

**Figure S1.**
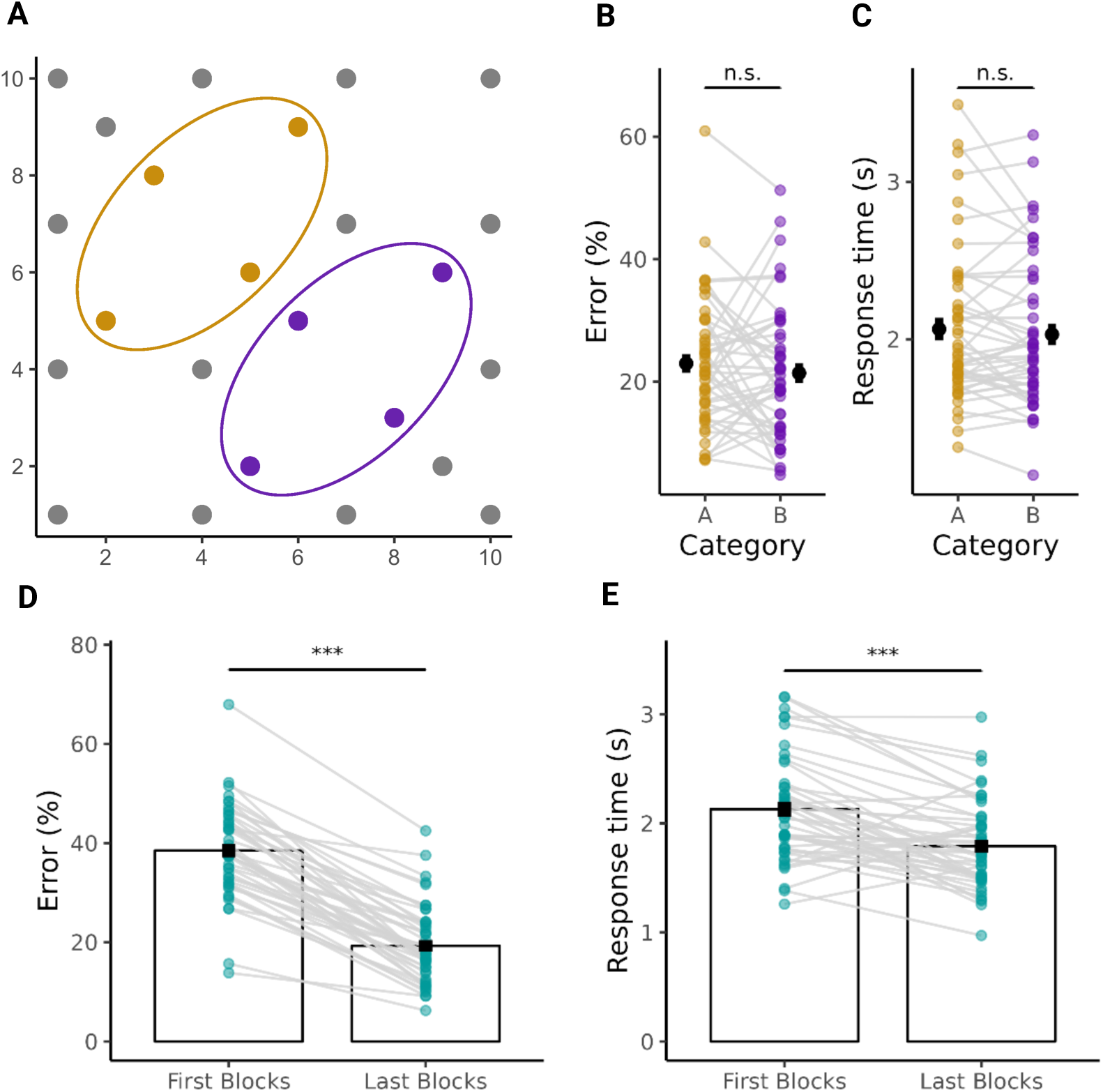
Results of the categorization training. **A:** Categorization stimuli (dots) were drawn from a two-dimensional feature space that embedded two elliptical-shaped categories (yellow & purple) and a residual category (gray). **B** & **C:** There was no significant (n.s.) difference between the two elliptical-shaped categories in categorization errors (*t*_46_ = -1.020, *P* = .313) and response time (*t*_46_ = -1.002, *P* = .322). Colored dots depict participant data points and black dots with error bars reflect the means +- SEM. **D** & **E:** Categorization performance improved between the first and the last five blocks, as indicated by a lower percentage of categorization errors (*t* = -18.164, *P* < .0001) and faster response time (*t* = -5.600, *P* < .0001). Green dots depict participants’ data points, and bars with error bars reflect the mean +- SEM. *** *P* < .001

**Figure S2.**
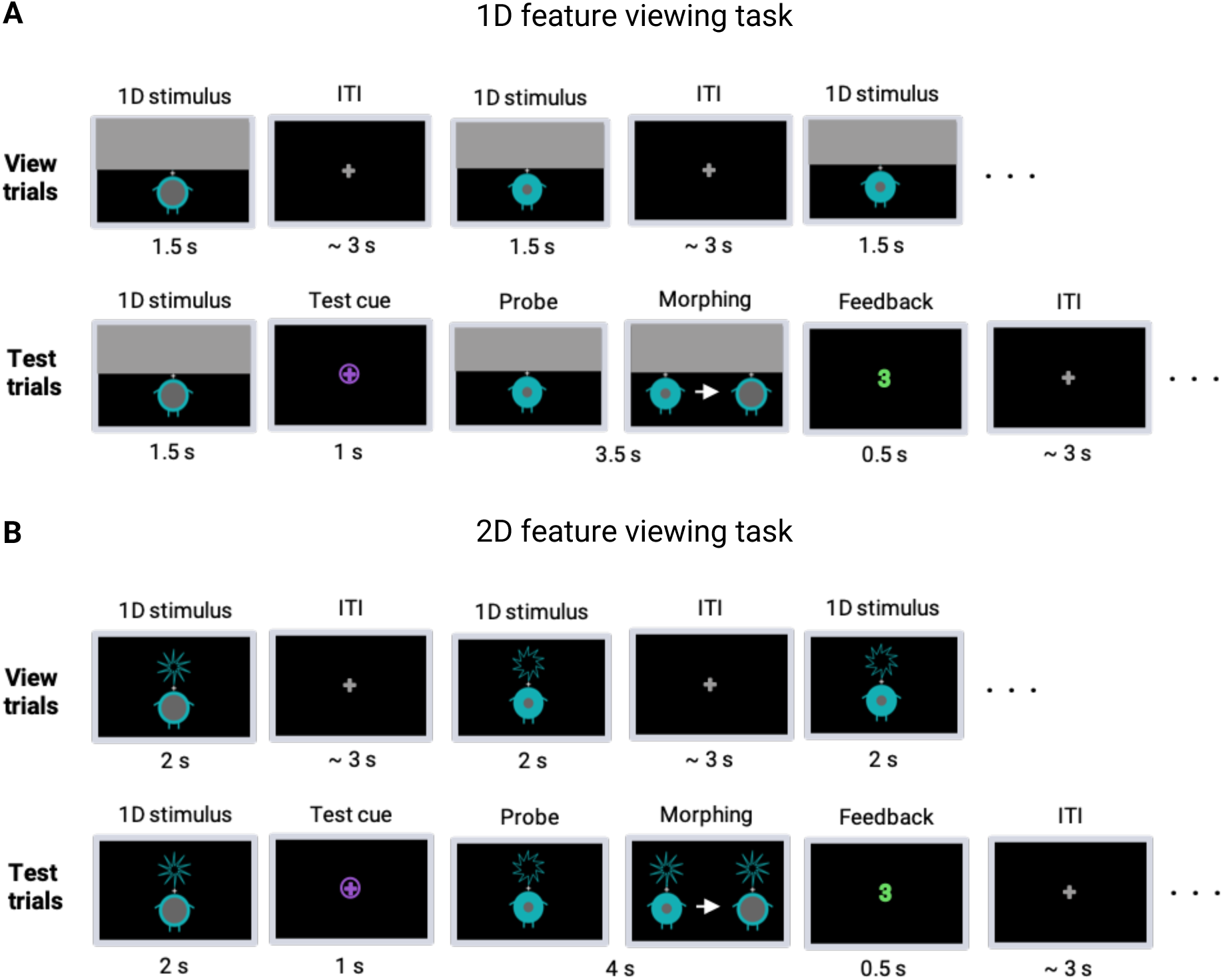
Trial display in the feature viewing and reconstruction tasks. **A:** In the 1D viewing task, participants were instructed to attend to one feature (here the stomach) of the stimuli. The other feature was occluded by a gray rectangle throughout the task. Stimuli were presented for 1.5 s and separated by a fixation cross (jittered intertrial intervals, mean 3 s). In addition to regular viewing trials, test trials (23%) were included to ensure attention to the feature values. Test trials were indicated by a purple cue (1 s). Participants’ task was to change the feature value presented after the purple cue into the feature value presented before the purple cue (via two buttons to increase/decrease the value). After confirming their choice, participants received a green-colored number as feedback (0.5 s) that indicated the distance between their provided and the correct response. **B:** The 2D viewing task followed the same procedure, except that complete 2D stimuli were shown without occlusion. In test trials, one of the two features had to be changed (here stomach) in order to reconstruct the previous stimulus.

**Figure S3.**
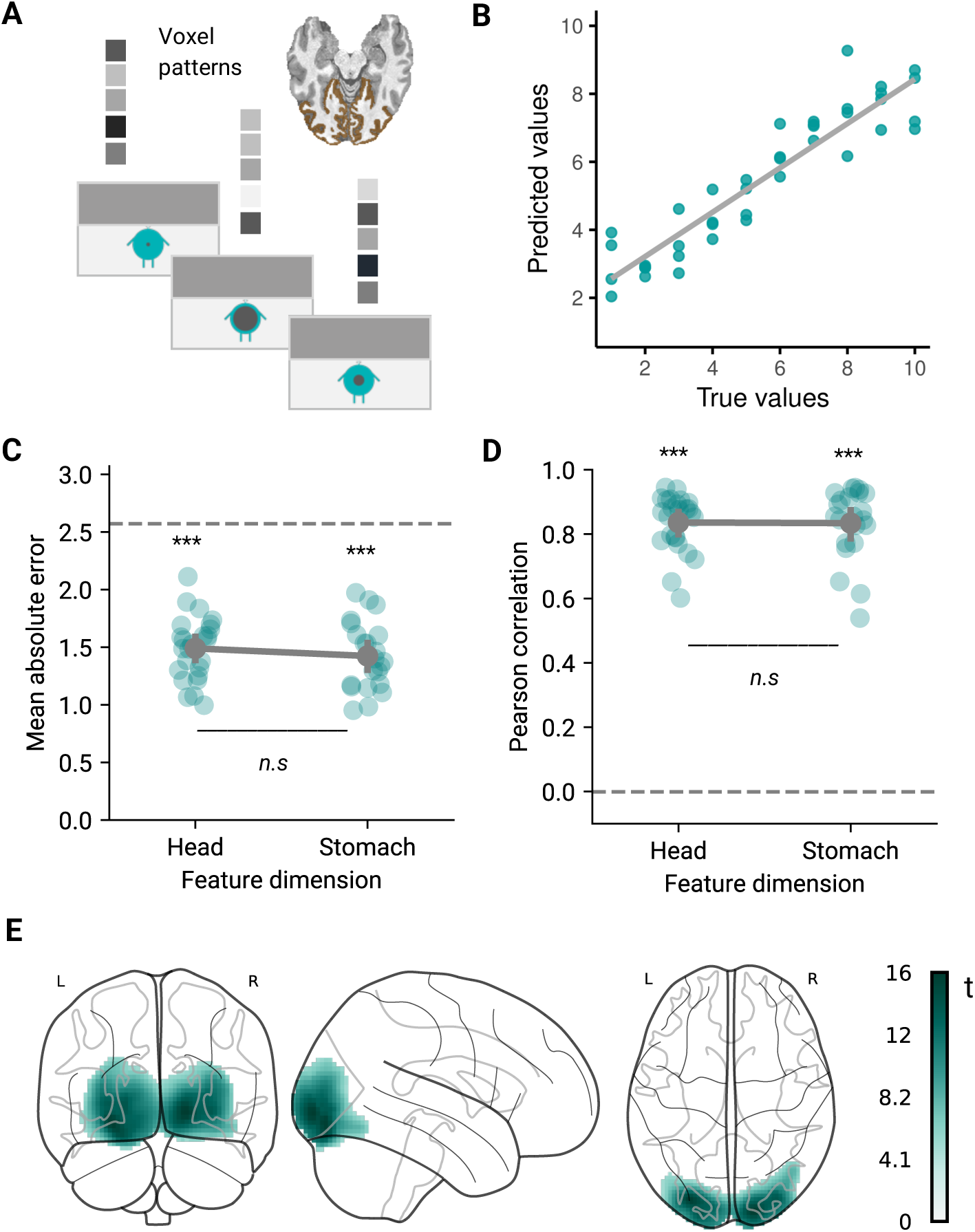
Training and validation of feature decoding. Related to Figure 4. **A:** A continuous decoder (linear support vector regression) was trained on multi-voxel patterns in visual cortex, reflecting run-wise GLM parameter estimates of different feature values. **B:** Decoding performance was evaluated in a leave-one-run-out (4-fold) cross-validation on the training data (depicted are four predicted values per feature value for an example participant with the regression line (gray)). **C:** Decoding performance was assessed via the deviation (mean absolute error) between true and predicted values. The scores were above chance level (*** = *P* < .0001) and did not differ between both dimensions (n.s., two-sample t-test: *t* = .07, *P* = .945). **D:** The same pattern was observed in the Pearson correlation coefficients between predicted and true values (*** = *P* < .0001; difference between dimensions: *t* = .79, *P* = .431). Green dots in C and D depict participant data points per condition; the gray dot and error bars refer to the mean value +- SEM. **E:** A whole-brain analysis confirmed that feature-predictive voxels (*R^2^*between predicted and true values) were constrained to visual cortex regions. Depicted are t-values from supra-threshold clusters on a glass brain (one-sided non-parametric permutation test with TFCE and *P*_FWE_ < 0.05).

**Table S1.**
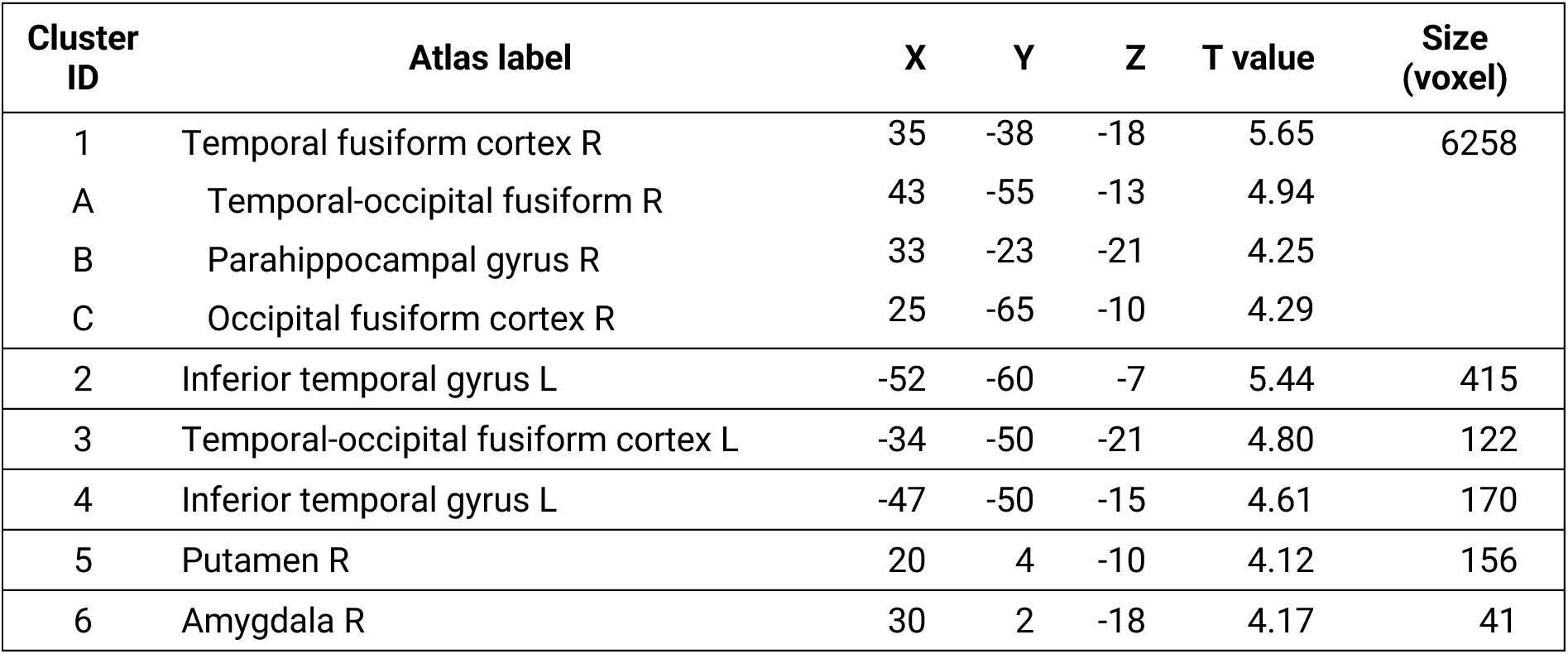
Representation of probe stimuli’s 2D-distance to the prototype examined via fMRI adaptation (related to Figure 5). An exploratory whole-brain analysis of the GLM described in Figure 5B further revealed significant clusters in the regions listed above (*P*_FWE_ < .05, TFCE; whole-brain corrected). Listed are atlas labels, MNI coordinates (X, Y, Z) and statistical T values of peak voxels from the clusters. Atlas labels are derived from the Harvard-Oxford (Sub)Cortical Structural Atlas. Sub-clusters are denoted by letters.

